# μECoG Recordings Through a Thinned Skull

**DOI:** 10.1101/564146

**Authors:** Sarah K. Brodnick, Jared P. Ness, Thomas J. Richner, Sanitta Thongpang, Joseph Novello, Mohammed Hayat, Kevin P Cheng, Lisa Krugner-Higby, Aaron J. Suminski, Kip A. Ludwig, Justin C. Williams

## Abstract

The studies described in this paper for the first time characterize the acute and chronic performance of optically transparent thin-film μECoG grids implanted on a thinned skull as both an electrophysiological complement to existing thinned skull preparation for optical recordings/manipulations, and a less invasive alternative to epidural or subdurally placed μECoG arrays. In a longitudinal chronic study, μECoG grids placed on top of a thinned skull maintain impedances comparable to epidurally placed μECoG grids that are stable for periods of at least one month. Optogenetic activation of cortex is also reliably demonstrated through the optically transparent ECoG grids acutely placed on the thinned skull. Finally, spatially distinct electrophysiological recordings were evident on μECoG electrodes placed on a thinned skull separated by 500-750μm, as assessed by stimulation evoked responses using optogenetic activation of cortex as well as invasive and epidermal stimulation of the sciatic and median nerve at chronic time points. Neural signals were collected through a thinned skull in multiple species, demonstrating potential utility in neuroscience research applications such as *in vivo* imaging, optogenetics, calcium imaging, and neurovascular coupling.

## 1 Introduction

Electrophysiological recordings of brain activity using high density electrode arrays are a staple of neuroscience research and have become increasingly prevalent for the clinical diagnosis of epileptic seizure foci as well as the clinical deployment of Brain-Machine Interfaces (BMI) [1; 2; 3]. Traditional electrophysiological recording methods involve the implantation of invasive electrode arrays either indwelling within cortex [4; 5], beneath the dura (subdural) [6; 7; 8], on top of the dura (epidural) [9; 10; 11], or non-invasively on the skin directly on above the exterior of the skull [12]. It is generally accepted that electrode placement closer to the neural signal sources of interest within the brain yields a more information rich and spatially distinct signal [13], whereas activity measured at a distance non-invasively is attenuated in part by the high impedance skull, yielding less spatially distinct information in the recorded signal from electrode to electrode [14; 15].

More recently, there has been a growing appreciation that surgical methods to open the skull, and/or the placement of an indwelling electrode grid on or within cortex, may cause adverse effects that impact the neural circuitry of interest [13; 16; 17]. For example, increased glial scarring [15; 18; 19], large increases in temperature of cortex [20], changes in intracranial pressure [21], intracranial hemorrhage and/or physical depression of cortex [18; 21; 22], and bacterial infection [21] have all been linked to the surgical procedure and implantation of electrocorticography (ECoG) or indwelling cortical arrays. These adverse events cause subtle changes to the neural circuitry of interest that have been shown to cause long-lasting deficit in performance of fine motor tasks amongst other consequences[16].

Concurrently there has been growing interest in neuroscience experiments that thin the skull instead of opening the skull to perform experiments using optical methods to record and manipulate both neuronal and non-neuronal cells within the brain [23; 24; 25; 26]. Removal of the skull in rodents has been shown to create glial scarring in the area under the craniotomy (Yang 2010 Nature Protocols, Yan 2009 J. Neurosci). Unlike the outer compact layer of the skull which has low conductivity, the spongy bone of the skull closer to the brain is low impedance [27] and if thinned appropriately is optically transparent [28]. However, the performance of μECoG grids placed chronically on a thinned skull preparation has yet to be evaluated.

To address this gap, a series of acute and chronic studies was performed where the skull was thinned to a translucent layer and implanted with a μECoG array. μECoG arrays were used in the experiments described because of their flexibility, transparency, and well characterized epidural signal profile [9; 10]. In rats chronically implanted for one month, impedance values and evoked somatosensory evoked potentials (SSEPs) were recorded at regular intervals to assess stability of electrical function and spatial resolution of recordings through the thinned skull. Cortical signals from optogenetic stimulation in a ChR2 mouse were recorded in an acute terminal session through a thinned skull and were compared to recordings through the dura after removal of the thinned skull. These studies tested multiple common stimulation paradigms for neuroscience research in multiple species to characterize the reliability and spatial resolution of electrophysiological recordings through a thinned skull.

## 2 Materials and Methods

### 2.1 Ethics Statement

All animal procedures were approved by the Institutional Animal Care and Use Committee (IACUC) at the University of Wisconsin – Madison. All efforts were made to minimize animal discomfort.

### 2.2 Device Fabrication

μECoG devices were fabricated following protocols previously described for polyimide [10] and Parylene C [29] arrays (Figure 1). Rat sized polyimide (750 um spacing, 250 um site diameter) (Figure 1b) or Parylene C (750 um spacing, 200 um site diameter) (Figure 1c) based μECoG electrode arrays were custom fabricated with 16 platinum sites (one or two 4mm × 4mm grids) and implanted unilaterally or bilaterally (Figure 1c) between bregma and lambda in Sprague-Dawley rats. Similarly, for experiments with mice, a smaller, 2mm × 2mm 16 platinum site Parylene-C μECoG array (500 um spacing, 150 um site diameter), was fabricated and used for optogenetic experiments (Figure 2c). Parylene C was chosen for optogenetic and imaging studies due to its flexibility and translucent properties.

**Figure 1:**
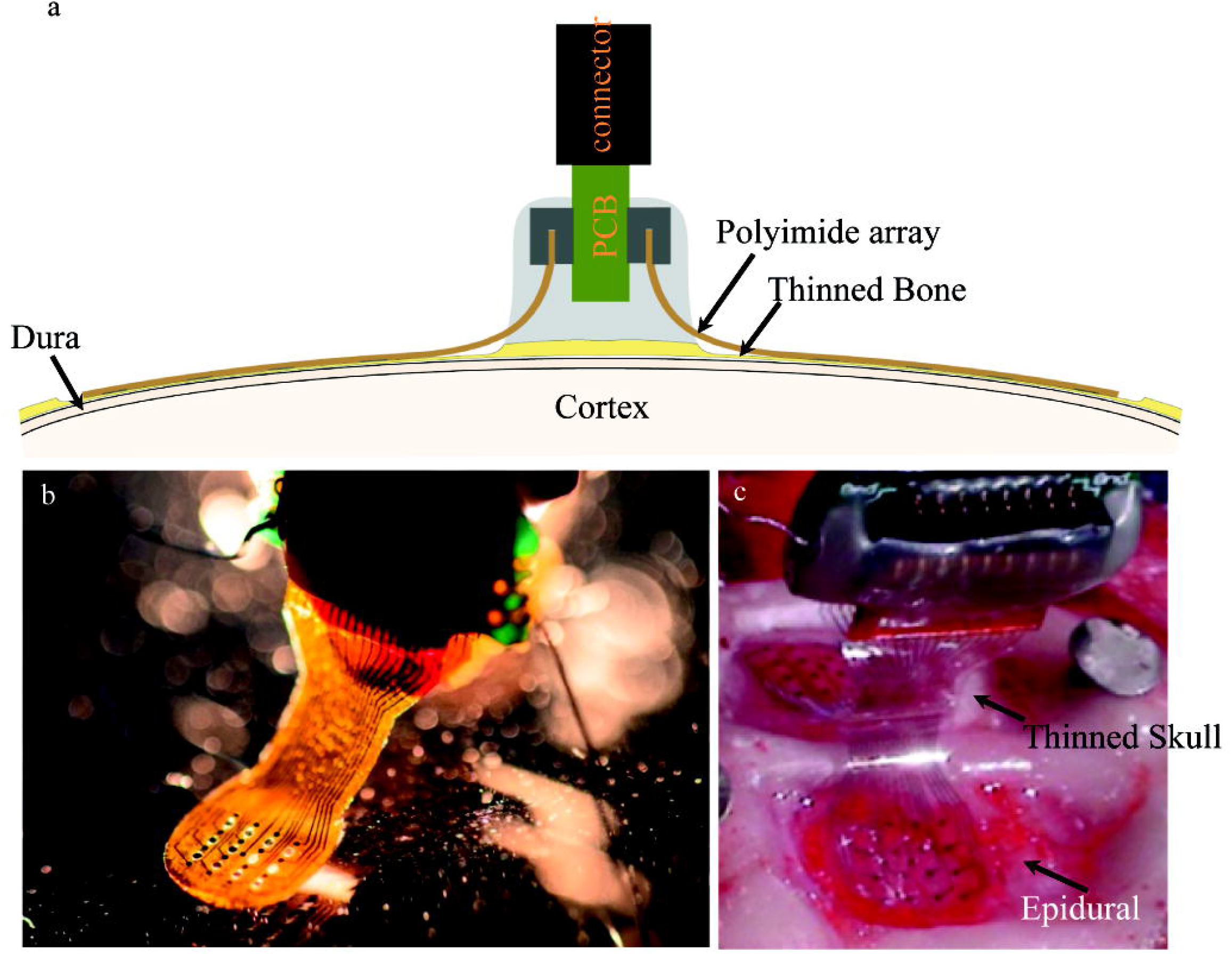
**(a)** Diagram illustrating chronic placement of 16-channel bilateral μECoG array, and ZIF connector on thinned skull surface over sensorimotor cortex in a rat. **(b)** Polyimide based platinum μECoG array with 16 channels (750 μm spacing, 250 μm site diameter). **(c)** Surgical photograph of bilateral Parylene C based platinum μECoG array being placed over a thinned skull (top of photograph) and on the dural surface (bottom of photograph).

**Figure 2:**
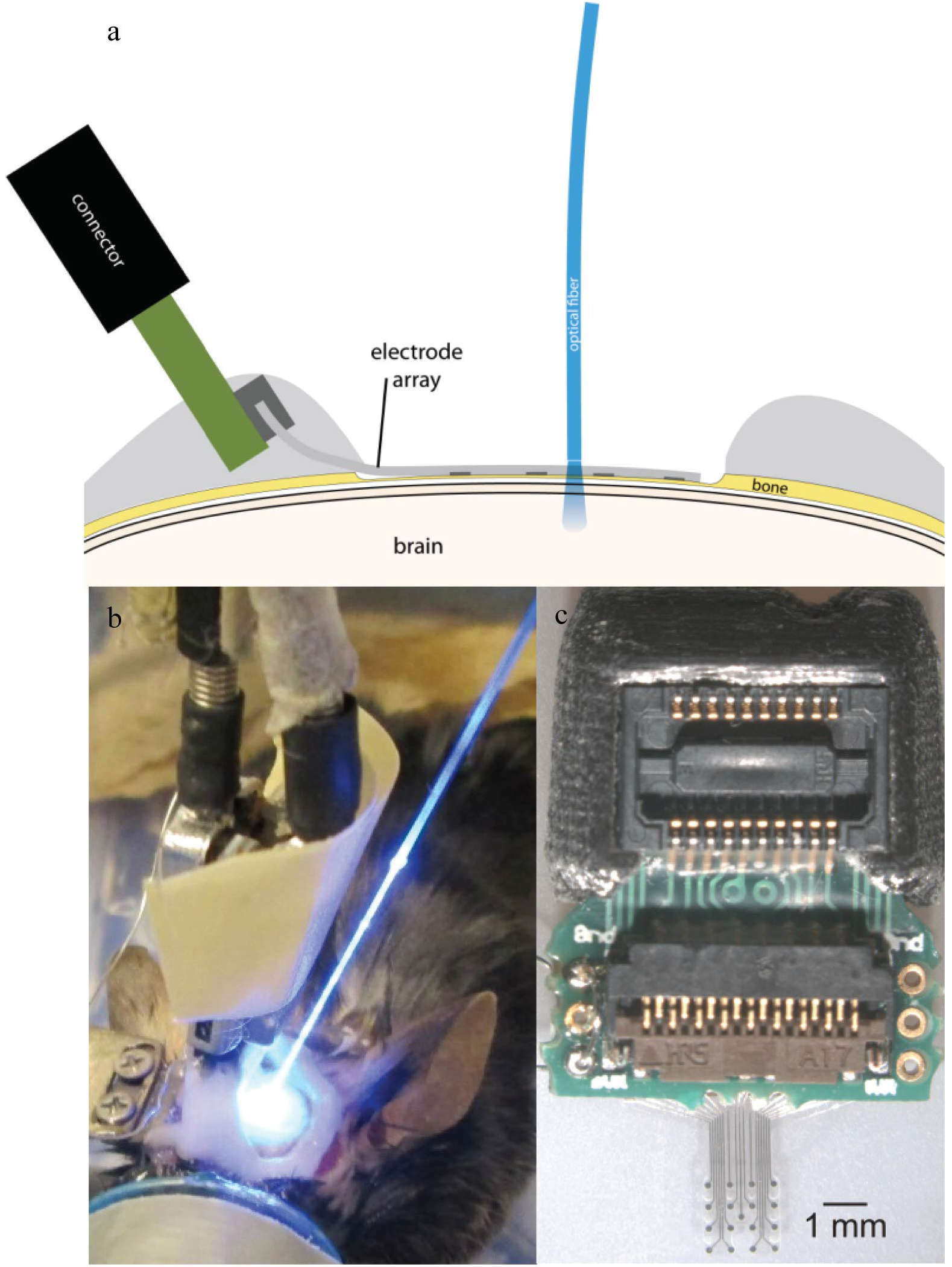
(a) Illustration of μECoG electrode array placement over a thinned skull in a mouse with optical fiber positioning. (b) Optogenetic stimulation of cortex with optical fiber placed on a μECoG array over either thinned skull. (c) Parylene C based platinum μECoG array with 16 channels (500 μm spacing, 150 μm site diameter) and ZIF connector.

### 2.3 Surgical Preparation

#### Chronic Experiments

Male Sprague-Dawley rats (n=7, Envigo, Indianapolis IN) 2-4 months old were chronically implanted with custom built μECoG arrays (Figure 1a). Three rats were implanted with bilateral arrays over thinned skull, three rats were implanted with bilateral arrays over the dural surface, and one rat was implanted with a bilateral array, one over thinned skull and one over the dura. The electrode array on the dural hemisphere for this animal was damaged during insertion and not viable therefore electrophysiology and impedance data were not included in this paper, however, histological staining was performed and included. Surgical procedures were based on previously published methods [30]. Before surgery, buprenorphine hydrochloride (0.05 mg kg^−1^, Reckitt Benckiser Healthcare) was administered for analgesia and dexamethasone (2 mg kg^−1^, AgriLabs) to prevent cerebral edema. Rats were induced with 5% isoflurane gas in O2 and maintained on 1.0-2.5% throughout the duration of the surgery. Following induction, rats were placed into a stereotaxic frame with the scalp shaved and prepped with alternating povidone iodine and alcohol. The skin was incised, and the exposed skull was cleaned and dried. Three stainless steel screws (stainless steel, 00–80 × 1/8 in.), two for attachment of a ground wire, and another for reference and mechanical support, were attached to the rostral and caudal areas of the skull. Next, UV curable dental acrylic (Fusio, Pentron Clinical) was placed on the periphery of the exposed skull to provide an anchor for the attachment of future acrylic, and two craniotomies (~5mm × 5mm) or thin skull areas were drilled over somatosensory cortex. A thinned skull area was made by drilling through the top layer of compact bone, through the spongy layer, and slightly into the lower compact bone where it became transparent. The μECoG arrays were placed epidurally or over a thinned skull area and covered with a thin layer of GelFoam (Pharmacia and Upjohn Co, New York, NY) and saline before being covered by dental acrylic. GelFoam was used to prevent acrylic from covering the electrode array and was only placed on top of the arrays and not beneath. The ZIF connector was then secured to the skull and a purse string suture (3-0 vicryl) closed the skin wound. Triple antibiotic ointment was applied to the wound during closing to prevent infection. Rats were monitored post-surgically until they were ambulatory and showed no signs of pain or distress. Another dose of buprenorphine was administered 8-12 hours after the initial dose to relieve any pain the animal may have been experiencing following the surgery. Ampicillin (50 mg kg^−1^ SC, Sage Pharmaceutical) was administered twice daily for seven days postoperatively to prevent infection.

### 2.4 Periodic Chronic Electrophysiology Testing

Sensorimotor evoked potentials were recorded periodically for up to one month under sedation in rats with chronic μECoG implants to assess signal stability and uniqueness/spatial resolution of information recorded on nearby sites. Dexmedetomidine (50 μgrams/kg subcutaneous injection (SC)) was used to achieve sedation. Atipamezole (0.5 mg/kg SC) was administered at the end of the procedure as a reversal agent. Dexmedetomidine sedation was supplemented with small amounts of isoflurane (0-0.5%) throughout the procedure to deepen sedation. The sciatic or median nerve, hindlimb or forelimb respectively, were stimulated weekly to evoke somatosensory evoked potentials (SSEPs). Needle or surface stimulation electrodes were used. Needle electrodes were placed on either side of the sciatic nerve, 3 mm apart. Surface electrodes were placed on shaved skin above the sciatic or median nerve, with a reference electrode placed below the leg. Stimulation pulses were applied using needle electrodes (monophasic 0-0.8mA for 2ms), or surface electrodes (monophasic 0.5-3.5mA for 1ms) both at approximately 0.5 Hz. The cortical response was recorded simultaneously at 24KHz using a multichannel recording system (TDT, Alachua, Florida).

### 2.5 Acute Terminal Experiments

Three Thy1::ChR2/H134R-YFP (ChR2) mice (Jackson Laboratory; stock number 012350) ~6-16 weeks old were implanted during acute terminal recording sessions with μECoG arrays implanted over the dura or a thinned skull area to compare neural signals recorded from light stimulation (Figure 2a, 2b). Evoked potential data from optogenetic stimulation were collected from three mice and strength duration curve data were collected from one mouse. Arrays were placed onto a thinned skull first, and then placed epidurally after removing the thinned skull. Mouse surgical procedures were similar to previously published methods [31]. Briefly, mice were administered buprenorphine hydrochloride (0.05 mg kg^−1^) and dexamathasone (1 mg kg^−1^ SC) before induction, induced, and maintained with 1-2.5% isoflurane. The animal was placed in a stereotaxic-like frame and a craniotomy or thinned skull was performed. A μECoG array was placed on the dura or thinned skull and ground and reference wires were coiled and placed on a small area of thinned skull on the contralateral hemisphere. GelFoam was not used in optogenetic studies. Instead the cortical surface was continually kept wet with a saline drip.

Heart rate and blood oxygen concentration in both species were monitored throughout the surgery using a pulse oximeter. Body temperature was monitored with a digital thermometer and regulated with a water-circulated heating blanket.

### 2.6 Acute Terminal Optogenetic Electrophysiological Testing

Optogenetically evoked potentials were recorded during a terminal procedure by shining light though a fiber coupled LASER system or LED through an optically transparent parylene μECoG onto the dura or thinned skull of ChR2 mice using previously reported methods (15) (figure 2). Photostimulation was accomplished by using an optical fiber (200μm in diameter, 0.22 NA, flat cleaved and polished, Thorlabs, Newton, NJ) connected to a 100mW 473 nm LASER (Laserglow, Toronto, ON) and controlled by a multichannel system (TDT, Alachua, FL). 2.5ms pulses, varying power settings, and random interstimulus intervals were used. Power at the tip of the optical fiber was approximately 80 mW/mm^2^ and was placed approximately 1mm from the cortical or thinned skull surface. Recordings were taken with a Tucker-Davis Technologies RZ2 amplifier (TDT, Alachua, FL), and sampled with a high impedance headstage. A photostimulus delivered by an LED (465 nm, RGB MC-E, Cree, Durham, NC) approximately 2 cm away from the cortical or thinned skull surface was used to create photostimulus duration versus amplitude peak to peak potential contour plots (figure 8). Voltage pulses were changed to current pulses (0–1000 mA, 0.5–12 ms) with an LED driver (BuckBlock, LEDdynamics, Randolph, VT). Irradiance was calculated by measuring optical power (PM100D, S130C, Thor Labs, Newton, NJ) 2 cm from the LED, and the result was divided by the commercially available photo sensor’s area (S370 Optometer, United Detector Technology, Hawthorne, CA).

### 2.7 Electrophysiology Analysis

Chronic electrical evoked responses were recorded through a thinned skull preparation. Optogenetically evoked responses from terminal mice experiments were recorded both epidurally and through a thin layer of skull. Local field potentials (LFPs) were bandpass filtered using a combination of a 2nd order, Butterworth lowpass filter (cutoff frequency = 1000 Hz), a Butterworth high pass filter (cutoff frequency = 3Hz), and a 3rd order notch filter (cutoff frequencies =55Hz and 65Hz) to remove line noise. Evoked potentials were computed from the average of evoked responses from the same stimulus amplitude and channel. To increase signal-to-noise ratios, two known post processing referencing techniques, common average referencing (CAR) and small Laplacian referencing, were employed and compared (Figure 3). Each were incorporated as described in the literature [32; 33]. After small Laplacian referencing, heatmaps of the electrical and optogenetic evoked responses (Figures 4 and 7), were created using the maximum positive and negative peaks to visualize the spatial organization of the cortical responses. Each peak was defined as the average of 7 data points centered at the maximum and minimum points of the response in a window defined as roughly 10ms to 35 ms post stimulus onset and subtracted [34; 35]. Channels above 600kOhms were considered to be non-functional and removed from analysis.

**Figure 3:**
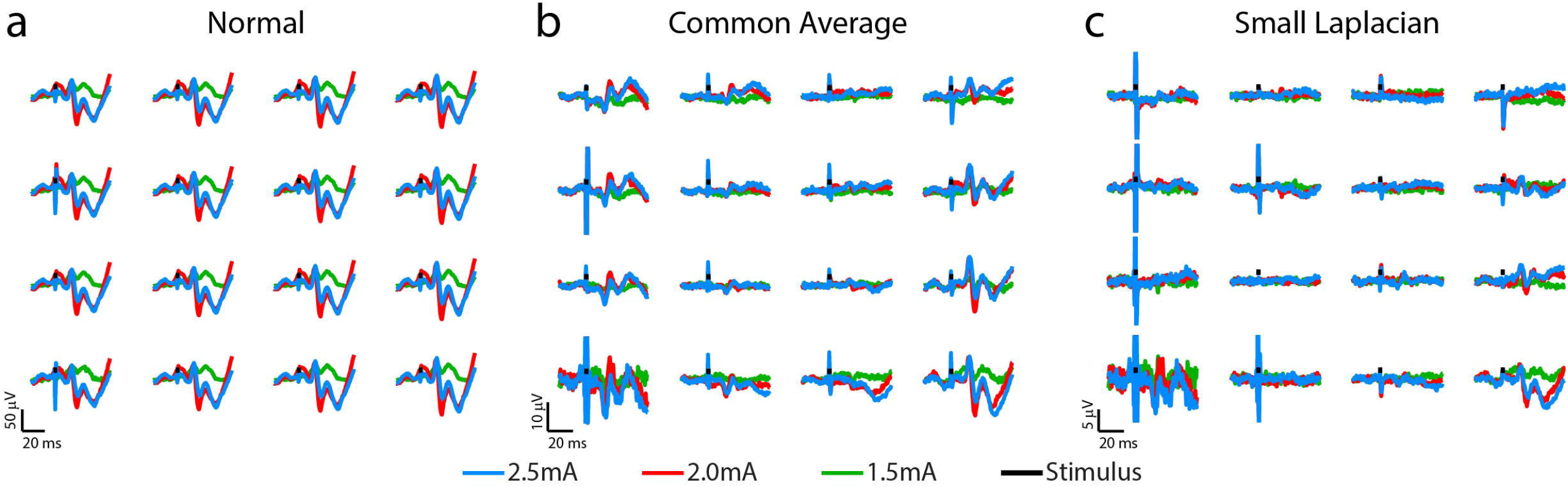
Somatosensory evoked potentials (SSEPs) recorded on day 20 post-implantation from a rat implanted with a 16-channel μECoG array placed over thinned skull left sensorimotor cortex. Biphasic current pulses (1 ms, varied amplitude) were used to stimulate the right hindlimb with surface electrodes over the sciatic nerve. (a) Stainless-steel bone screw, (b) Common Average and (c) small Laplacian referencing strategies are shown to increase the signal-to-noise ratio, and to reveal spatial signaling from the predicted hindlimb anatomical region.

**Figure 4:**
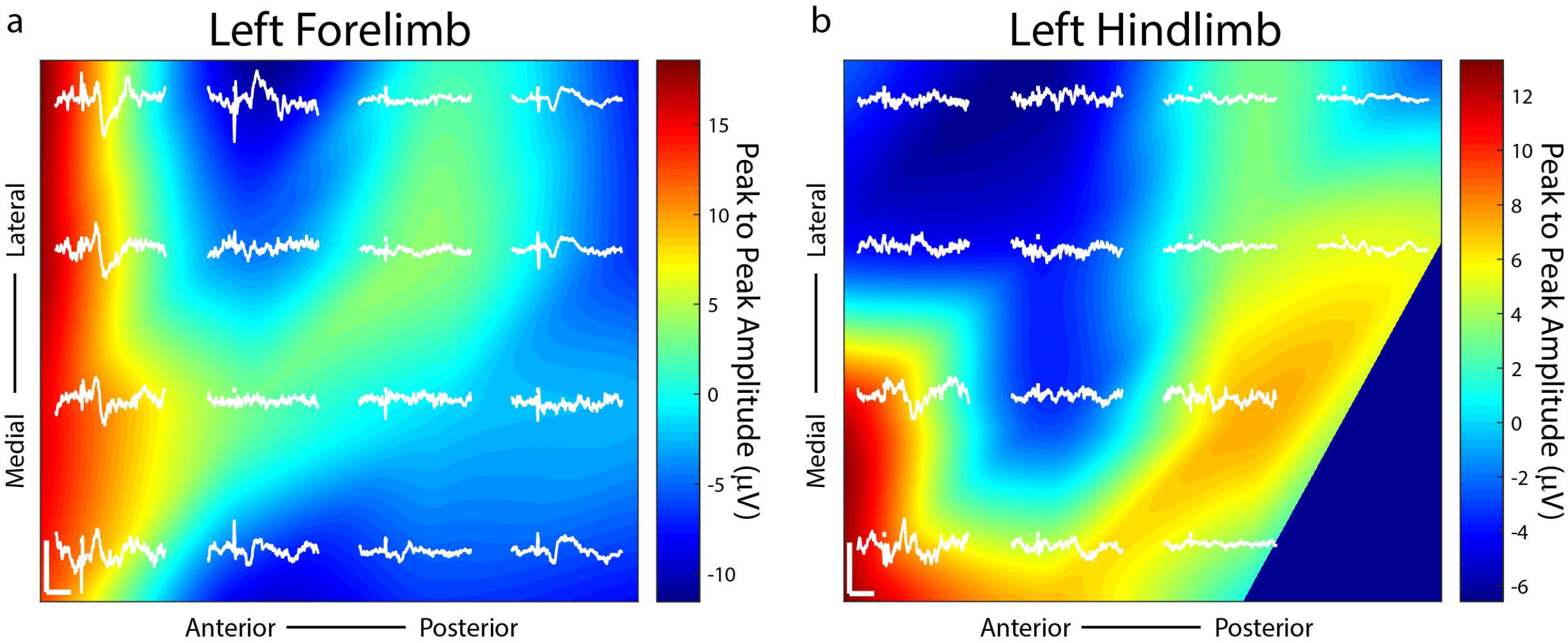
Somatosensory evoked potentials (SSEPs) on day 38 post-implantation with small Laplacian referencing from forelimb and hindlimb electrical surface stimulation using a 16-channel μECoG array placed over a thinned skull portion of rat sensorimotor cortex. Plots represent spatial recordings from the same electrode array, demonstrating LFPs from (a) biphasic forelimb stimulation and (b) monophasic hindlimb stimulation. Stimuli were applied for 1ms at 1.25mA. Activity is represented by 2D interpolated heat maps. The portions closer to the red spectrum show evoked activity higher than baseline when averaged over at least 25 trials, and closer to blue shows negative activity. The *x* scale bar, 20 ms; *y* scale bar, 20 μV.

### 2.8 Chronic Impedance Recordings and Analysis

Electrical impedance spectra were collected from arrays before implantation, and periodically after implantation to assess electrical characteristics using a potentiostat (Autolab PGSTAT 128N, Metrohm, Riverview, FL) and following previously published methods (Figure 5) [36]. Arrays that were determined viable for implantation had values of approximately 50-100 kΩ at 1 kHz. Animals were trained to sit still with treats and were not anesthetized or sedated for chronic impedance measurements. Analysis consisted of data from six rats, three with thinned skull implants and three with epidural implants for comparison. Each rat had a bilateral implant consisting of 32 electrode sites. Impedance measurements were gathered from each electrode for 30 days post implantation. Resistive values at 1kHz were plotted for each of the 32 channels corresponding to length of time of the implant. Single channels with a resistance above 600 kΩ were considered outliers and eliminated from calculations for that day. Outliers were considered to be broken or due to an inadequate connection. Average resistance was plotted for each day and fitted to a curve across days using cubic spline interpolation to account for measurements potentially not lining up exactly on individual days across animals. The thinned skull implants interpolation curves were averaged together and plotted against the epidural implants averaged interpolation curves. Impedance values were not recorded after implantation in acute mice experiments, although pre-implantation impedance values were comparable to those of the devices implanted for chronic recordings in rats.

**Figure 5:**
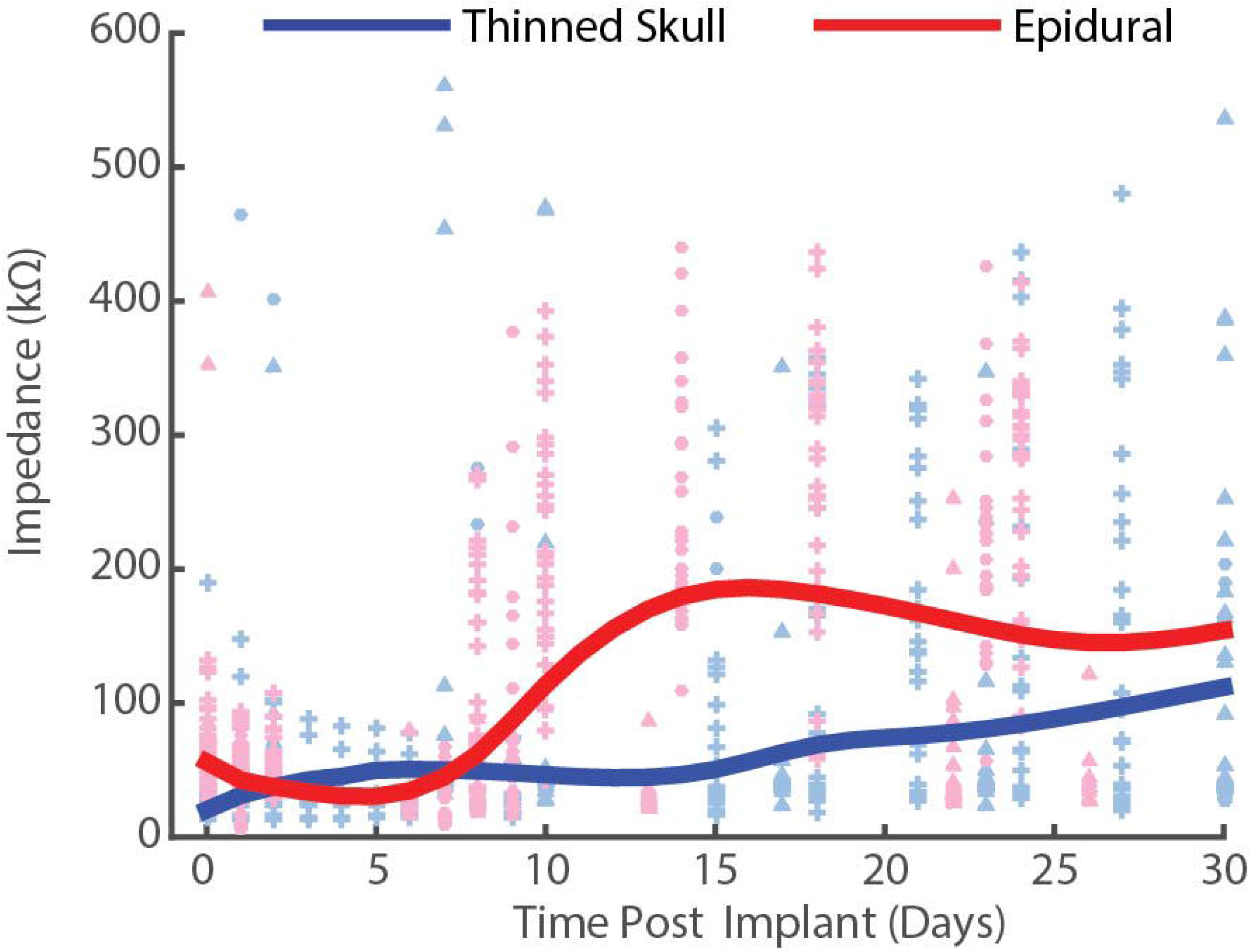
(a) Chronic impedance spectral data at 1kHz from thinned skull (blue) and epidurally (red) implanted μECoG electrodes in rats. Each interpolation curve represents three animals per group, and 32 electrode sites per animal. Individual data points represent individual electrode site impedance spectra measurements. Epidural impedances increase until approximately two weeks and plateau, while thinned skull impedances remain lower and more stable.

## 3.0 Results and Discussion

### 3.1 Chronic periodic sensory evoked potential recordings in rats

Chronic SSEP recordings were obtained weekly during electrical stimulation of the sciatic or median nerve in three rats to compare the spatial resolution of thinned skull μECoG arrays versus traditional epidural arrays. Thinned skull μECoG arrays were implanted bilaterally in sensorimotor cortex in each rat and SSEPs were recorded in each contralateral hemisphere from a cutaneous electrical stimulus of hindlimb or forelimb. A representative plot of thinned skull SSEPs on the left hemisphere from right hindlimb stimulation is shown in Figure 3. Using the stainless-steel bone screw as a reference, which was implanted cranial and contralateral to the μECoG array (Figure 3a), recorded signals contained common noise, and differences in SSEPs from nearby electrode locations were not readily apparent. Two post process referencing techniques were used to reduce both common noise and common signal to highlight spatially distinct differences in neural signals.

Employing a Common Average Reference (CAR) (Figure 3b) successfully recovered spatially distinct hindlimb SSEPs on adjacent electrode sites. Similarly, employing a small Laplacian (Figure 3c) reference post-hoc further highlighted spatially distinct SSEP responses on adjacent sites. Consequently, we chose to use the small Laplacian post-hoc referencing strategy for the remainder of the recording data, because it visually increased unique highlighted spatial information present in the SSEP on adjacent sites [32].

Distinct somatotopic signals were recorded 38 days post-implantation from μECoG arrays placed on a thinned skull area of the rat’s right sensorimotor cortex from both left hindlimb and forelimb stimulation (Figure 4). Small Laplacian referencing methods were also applied. Highest peak to peak SSEP values from forelimb stimulation, according to the heatmap, are positioned at the anterior portion of the electrode with peaks spanning both medially and laterally (Figure 4a). When switching the area of stimulation to the hindlimb, SSEPs shifted medially similar to previously mapped rat sensorimotor cortex (Figure 4b) [35]. The recorded sensory responses are consistent with the response latencies for myelinated sensory fiber conduction, around 13ms for forelimb and 17ms for hindlimb according to previously published data [34; 35].

Thinned skull μECoG electrode arrays not surprisingly have lower signal amplitudes recorded during evoked responses in comparison to historical studies using the same arrays placed epidurally which are closer to the source of the neural signal [9]. As a result, it becomes more important to employ CAR and small Laplacian referencing strategies to eliminate common signal/noise to uncover spatially distinct spatial information for neuroscience applications. Given a similar SSEP was recorded across all electrode sites with appropriate conduction latency prior to post-hoc referencing, this may suggest the common signal was recorded at the stainless-steel bone screws in contact with the surface of the brain used for the reference and ground respectively. Although the ECoG signal recorded from the bone screws has been insignificant compared to signals recorded epidurally from μECoG in previous studies and therefore post-hoc referencing was not required to reveal spatially distinct information from site to site, the attenuation of signal through the thin skull made the small common signal putatively recorded from the stainless-steel screws more problematic. Consequently, post-hoc referencing was necessary before spatially distinct SSEPs were observed on adjacent electrode sites.

Thinned skull μECoG electrode arrays also have been shown in this study to record information on a temporal scale similar to epidurally placed arrays. For example, thinned skull SSEPs were recorded at a temporal resolution of roughly 15ms. Currently, GCaMP6f is a popular genetically coded calcium indication (GECI) that is commonly used to observe neural activity at an onset of approximately 45ms [37]. This suggests that the incorporation of an optically transparent μECoG array with common thinned skull experiments for optical imaging would provide unique, complementary temporal information.

### 3.2 Chronic periodic impedance spectra recordings in rats

To compare the electrical performance of epidural vs thinned skull placed electrodes in rats over time, we measured the impedance spectra of electrodes on each array at 1kHz periodically over the chronic implantation period (Figure 5). Impedance plots from μECoG arrays implanted on a thinned skull preparation showed significantly different patterns of change over time than those implanted epidurally (Figure 5). Initial electrode impedances were similar when measured in 0.9 % w/v phosphate buffered NaCl saline (~25-125mOhms at 1KHz). After approximately 14 days of implantation as shown in Figure 5, the impedances of the electrodes on the epidural surface were higher on average than that of the electrodes on the thinned skull surface. The epidural impedance interpolation curve shows rise in impedance around one week after implantation and lasting for approximately 14 days, similar to other microelectrodes implanted in or on cortex in other studies [36; 38]. This may be attributed to a central nervous system immune response and new tissue formation and follows previous intracortical and epidural implantation impedance results [9; 36]. Impedances of epidural implants reached a steady state between 2-4 weeks post-implant reflecting decelerated wound healing. In contrast, the thinned skull impedance interpolation curve remained relatively stable for approximately 21 days post-surgery, only rising slightly towards the end of the 30-day period. This suggests the chronic electrode/tissue interface is different in composition than the epidural grids, but also demonstrates that bone regrowth/scarring under the thinned skull electrodes does not significantly increase the impedance by comparison. Supplementary figure 1 shows line plots of individually recorded impedance values from each rat over a time period of one month.

Decreased impedances during the first few weeks of thinned skull electrode implantation may suggest edema, and that fluid remained at the electrode/tissue interface without clearing. Extra fluid could have hypothetically caused shunting of current and increased distance between the electrode array and the thinned skull. Regardless, we were still able to record spatially and temporally accurate SSEPs and optogenetically induced filed potentials from thinned skull electrodes in rats and mice with relatively low impedance values (impedance values not acquired post-implantation in acute mice studies).

### 3.3 Comparison of thinned skull vs epidural recordings from optogenetic light stimulation in acute terminal mice

To further investigate the spatial resolution of information on nearby electrodes given a thinned skull recording approach, a light stimulus was applied through a clear Parylene C μECoG array and thinned skull to optogenetically activate neurons expressing light sensitive proteins. Optogenetically evoked potentials were recorded in ChR2 mice through a thinned skull (figure 6a) and epidurally using a smaller 2mm by 2mm clear μECoG array to generate a consistent focal activation of cortex for comparison. Evoked responses through the thinned skull showed the highest peak responses near the foci of optogenetic stimulation after small Laplacian referencing, further demonstrating spatiality the spatial recording ability of the preparation (Figure 6a). Increasing 473nm laser power also increased evoked potential peaks amplitudes. A similar response paradigm occurs with epidural stimulation (figure 6b), however, we obtain a much larger signal possibly due to lack of spatial filtration of signal through the skull. Figures 6c and 6d use a 2D interpolated heat map to show differences in peak to peak amplitudes at a stimulation laser power of 545.5mW/mm2. Both thinned skull and epidural heat maps display spatial distinct recordings on nearby electrode sites. Due to the presumed filtration/attenuation of signal through the skull and other tissues, the thinned skull recording (Figure 6c) is approximately 10 times less in peak-to-peak amplitude than the epidural recording (Figure 6d). The thinned skull signal also seems to be slightly more diffuse given appropriate referencing strategies.

**Figure 6:**
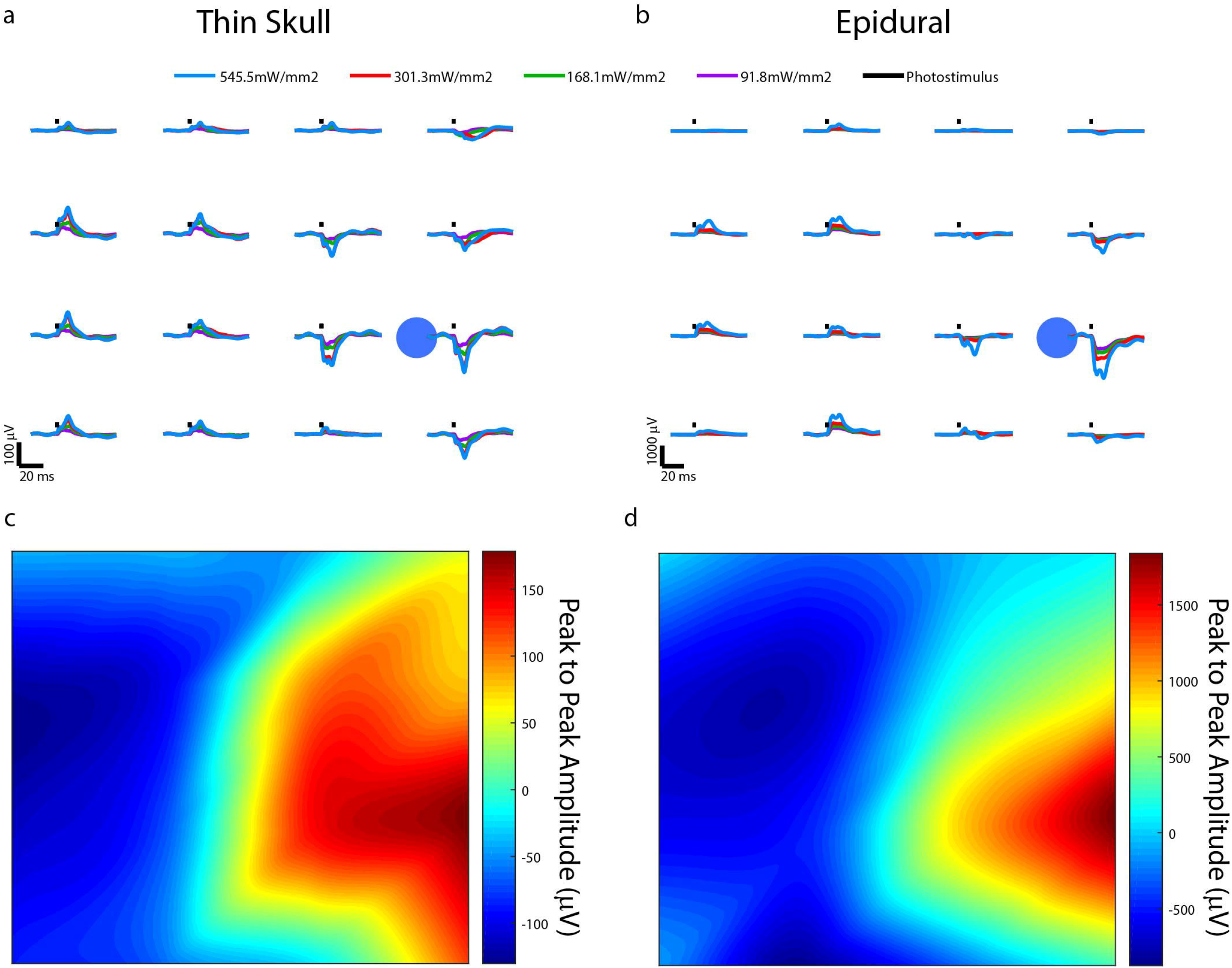
Optically evoked local field potentials from thinned skull (a) and epidurally (b) implanted μECoG arrays in an acute, terminal Thy1-ChR2 mouse. Amplitude heat maps show the 545.5 mW/mm2 optically evoked potentials using small Laplacian referencing from both the (c) thin skull and (d) epidural preparations. Each can be processed to illustrate the spatial resolution of the recordings, although the difference in scale is smaller in the thinned skull preparation by approximately a magnitude of 10.

One ChR2 mouse underwent an acute procedure where the skull was thinned, and a 465nm LED was positioned 2 cm away from the thinned skull with the light power and duration values varied to generate photostimulus strength vs duration curves (Figure 7). The resulting illumination covered most of the cortical area under the μECoG array. Stimuli strength and duration were applied randomly, and peak amplitudes of signals recorded. The thinned skull was then removed exposing the dural surface and stimulation procedure repeated. Figure 7a depicts a contour plots for signals recorded from the dura, whereas figure 7b depicts the same plot from the thinned skull.

**Figure 7:**
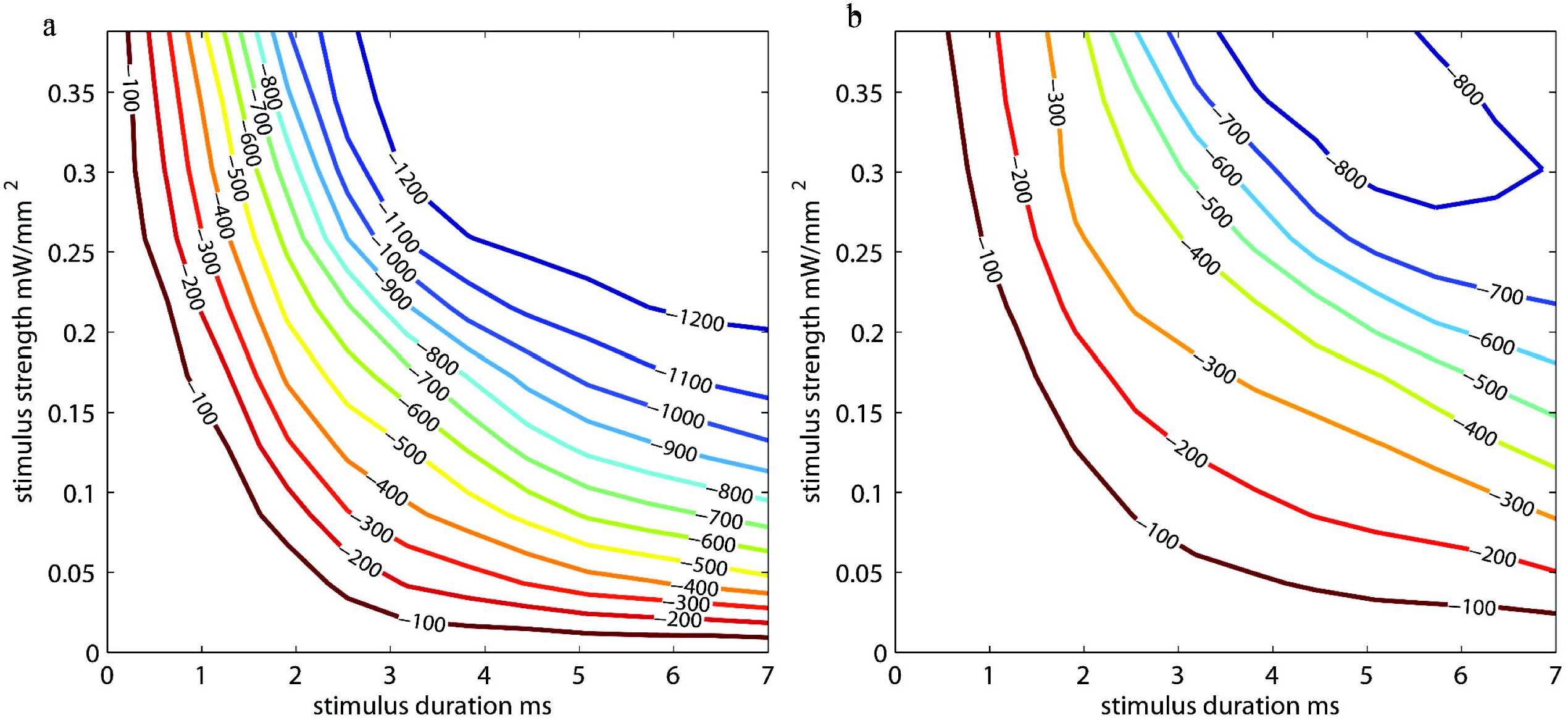
Photostimulus duration versus amplitude peak potential 2D interpolated contour plot. Stimulus strength is plotted against stimulus duration. Interpolated curves denoting the peak depolarization amplitudes (in μV) for the stimulus strength/duration are shown for (a) epidural and (b) thinned skull μECoG recordings in an acute terminal Thy1-ChR2 mouse. Longer stimulus durations and stimulus strength (power) are needed to evoke similar sized neural signal amplitudes in the thinned skull versus epidural preparations.

The optogenetically evoked μECoG signal on both the epidural and thinned skull grids demonstrated spatially distinct information, with waveform reversals often apparent on two adjacent sites. These reversals, in conjunction with the waveshape of the evoked response, demonstrated that the electrophysiological recordings were not photoelectric artifacts. Although the magnitude in μVolts of the evoked signal was approximately 10x less with the thinned skull preparation than with the epidurally placed grids, the spatial information as assessed by differences in recordings at adjacent electrode sites was highly similar after post-hoc Small Laplacian referencing.

### 3.4 Imaging of immune response in neural tissue to thinned skull and epidural μECoG implantations

Given the impedance responses over time in all animals, histology was performed on rat M32 to compare histology to the impedance measurement of approximately 50kOhms at timepoint 32 days post-implantation. Histologic sectioning and immunochemical staining for glial fibrillary acidic protein (GFAP) for astrocytes and Iba-1 for microglia and/or infiltrating macrophages was performed with perfused neural tissue in the single rat with bilateral thinned skull/epidural μECoG arrays. (Supplementary figure 2). The cortical region directly beneath the epidural preparation showed putative increase in GFAP immunoreactivity and projection of astrocytic processes towards the cortical surface (Supplementary figure 2c). Iba-1 staining did not reveal any obvious increases in microglial immunoreactivity within the brain tissue on either the epidural or thinned skull hemispheres (Supplementary figure 2d). However, an apparent thickening of the dura on the epidural side was observed (Supplementary figure 2f) which contained a higher density of Iba-1 positive cells, either microglia or infiltrating macrophages, that was not present on the thinned skull side of the animal (Supplementary figure 2e). Previous studies have also reported thickening of the dura under the μECoG array consisting primarily of collagen [22; 39].

Gross visual inspection of the brain surface post-mortem directly under the thinned skull where a μECoG array was chronically placed was also directly compared to the surface of the brain directly under a chronically implanted epidural μECoG array (see Supplementary Figure 3). Consistent with data reported by Onal 2003 [21] and Degenhart 2016 [22], the epidurally placed chronic μECoG array leaves a visible dent/impression on the surface of the brain upon removal. Similarly, denting is also visible beneath the two stainless steel mechanical and reference screws placed posterior to the olfactory bulbs. In contrast, no denting of the brain is evident under the thinned skull where the μECoG array was chronically placed.

The main benefit of the thinned skull preparation is that the skull remains partially intact. When the skull is completely removed many side effects can occur which may impact the interpretation of behavioral results. Previous studies in the field of *in vivo* imaging have showed increased glial reaction (microglia and astrocytes) under an open craniotomy window preparation compared to a thinned skull window preparation in mice [23]. Pneumocephalus can occur after craniotomy in a clinical setting which involves air being trapped in the cranial cavity [40]. Also, dendritic spine plasticity has been shown to differ in thinned versus open-skull window preparations, emphasizing that the neural environment under a craniotomy may be changed by the craniotomy itself [23; 41]. Another benefit the thinned skull recording technique might offer is improved implant mechanical stability, and the lack of direct contact with the surface of the brain or dura. The latter may reduce the risk of injury to neural tissue or device failure due to the lack of device movement on the surface of the brain, although this will need to be investigated in further studies.

## 4.0 Conclusion

In summary, the studies described in this paper for the first time demonstrate that μECoG grids placed on a thin skulled can provide stable, spatially distinct electrophysiological information out to periods in excess of a month. This method may be particularly useful as a complement to other studies which thin the skull for optical recordings/modulation of neural activity. μECoG recording grids may also provide a useful balance between invasiveness, information content, and day to day stability that may be important for neuroprosthetics applications. In addition, this novel method may be critically enabling for neuroscience studies in which minimizing the trauma to the underlying neural or non-neuronal cells of interest is necessary to avoid potential confounds given the fundamental hypothesis to be tested.

## 6.0 Author Contributions

SB, JN and TR: experimental design, data analysis, and manuscript preparation. AS, KL and JW: experimental design and manuscript preparation. KC, MH, and JN: data collection and data analysis. ST for device fabrication, and data collection. LKH contributed to manuscript preparation. All authors contributed to draft the manuscript and have read and approved the final manuscript.,

