## Supplementary figures and images for "μECoG Recordings Through a Thinned Skull"

### Supplementary Figure 1.

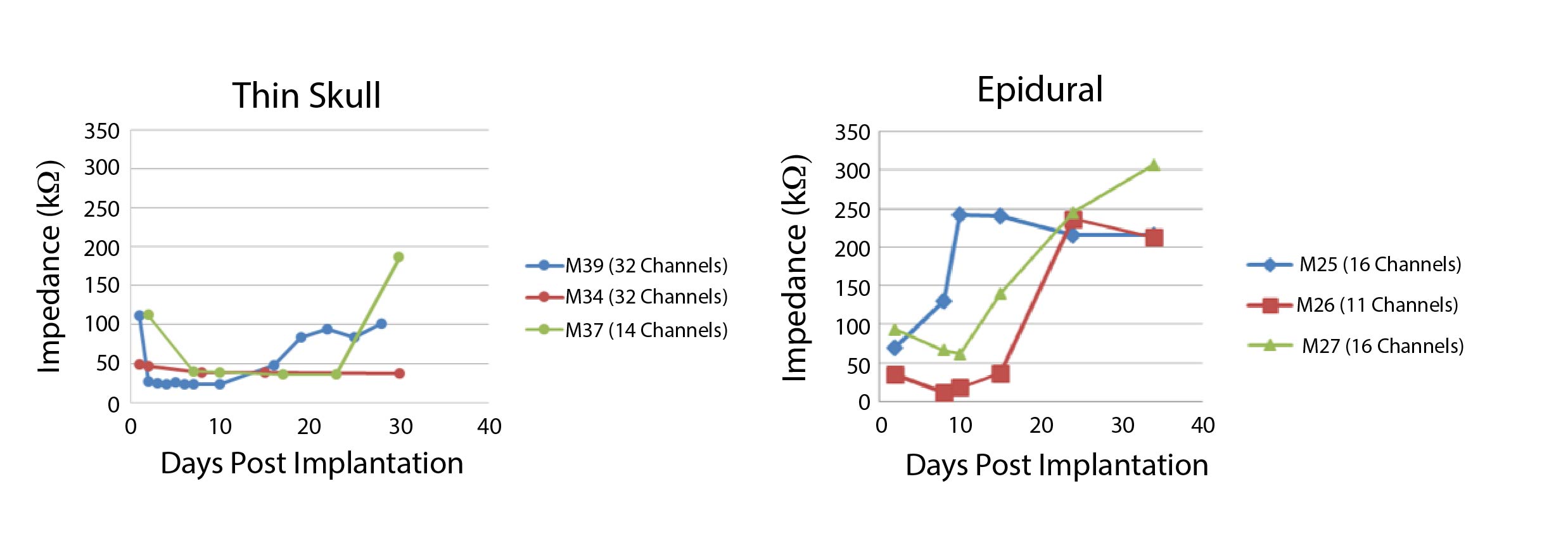

### Supplementary Figure 2.

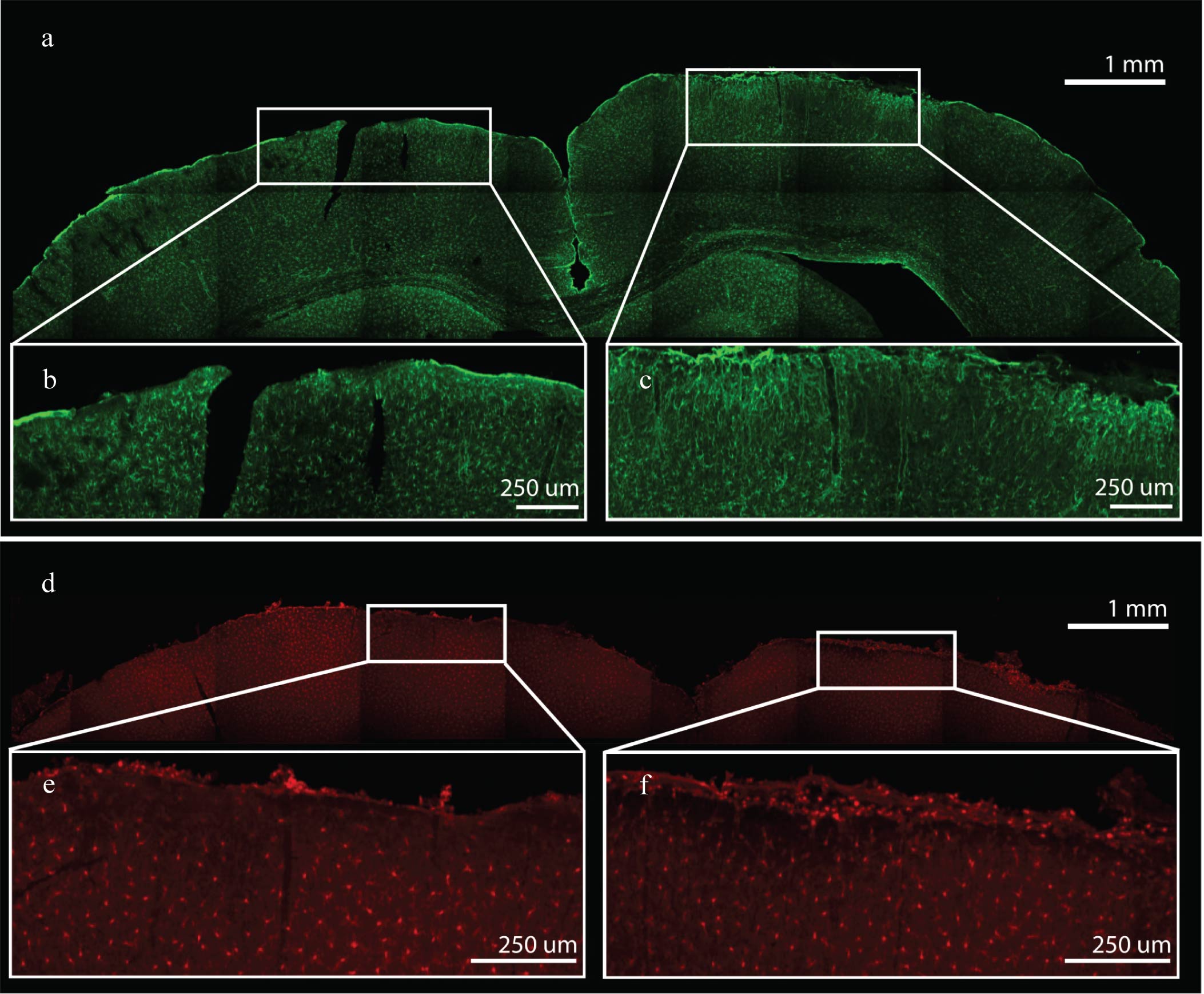

### Supplementary Figure 3.

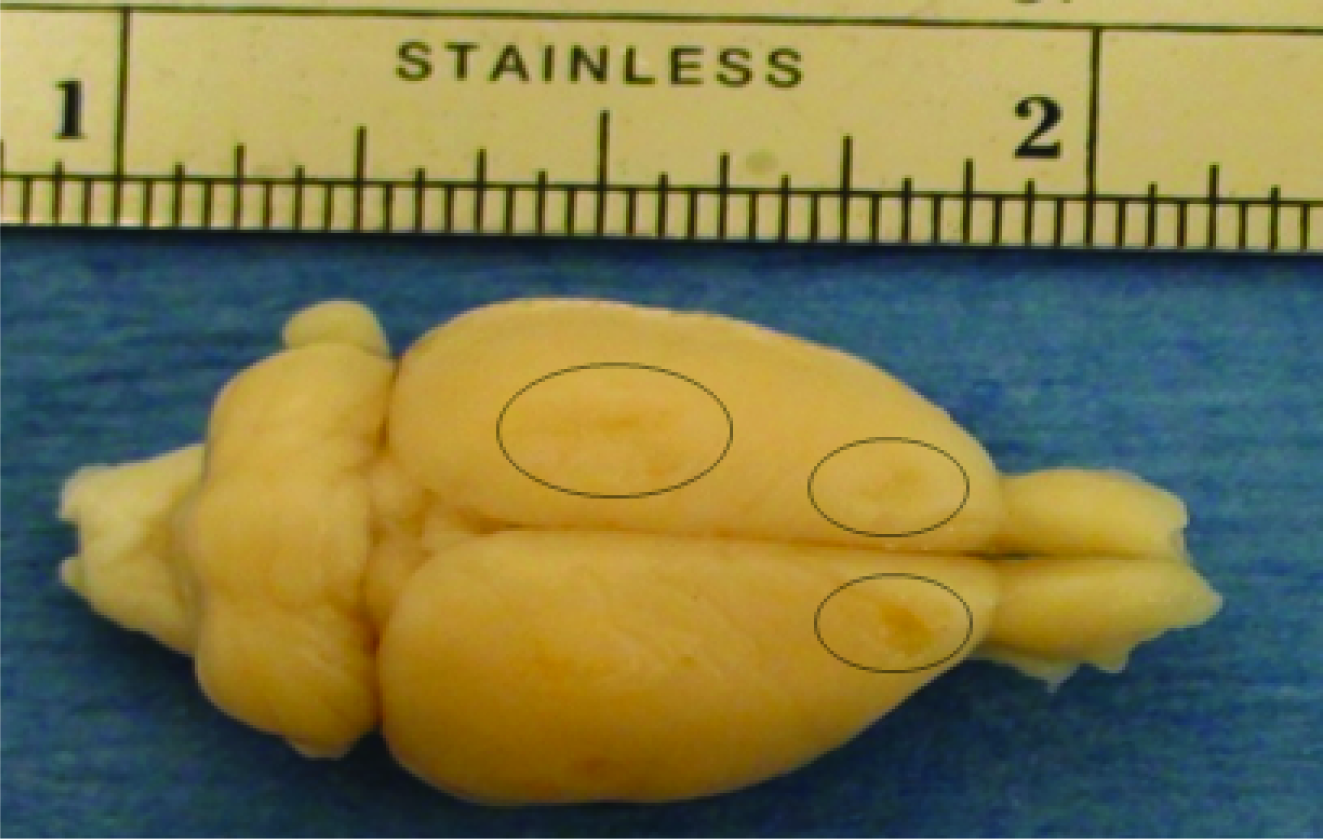
