## Supplementary Methods for "μECoG Recordings Through a Thinned Skull"

**Histology Methods**

Animals were deeply anesthetized using isoflurane and perfused with phosphate buffered saline (PBS) followed by a 4% paraformaldehyde (PFA) solution (Sigma-Aldrich) at the end of the experimental period. The whole brain was immediately extracted and left to soak in 4% PFA overnight after which it was moved to a staining buffer HEPES-buffered Hank's Balanced Salt Solution (HBSS) containing 90 mg/L sodium azide (Sigma-Aldrich). Coronal tissue slices 80 µm thick were cut on a Leica Vibratome (Leica Biosystems, Richmond, IL) from a region spanning several millimeters directly beneath the µECoG array. The slices were briefly incubated in a 5 mg/mL sodium borohydride (Sigma-Aldrich) solution to reduce PFA-derived autofluorescence followed by a permeabilization step using 0.5% Triton-X (ThermoFisher). Non-specific staining was blocked by overnight incubation in a 5% bovine serum albumin (Sigma-Aldrich) solution. The following day slices were stained overnight on glass coverslips using either mouse anti-GFAP (Sigma-Aldrich, cat. #3893) for astrocytes or rabbit anti-Iba1 (Wako, cat. #1919741) for microglia. After washing, secondary antibodies including goat anti-mouse AlexaFluor488 (Invitrogen, cat. #A-11034) and goat anti-rabbit AlexaFluor568 (Invitrogen, cat. #A-11031) were applied and the slices were mounted using ProLong Gold (Invitrogen, Carlsbad, CA). Slices were imaged on a custom-built multi-photon microscope with an automated XYZ stage and fitted with a 10X Nikon air objective and appropriate filter cubes. Images of the entire cortex were derived by programming the system to take sequential adjacent images that were combined using an ImageJ plug-in (Supplementary Figure 2).
